# One Hundred and Fifty Years of Warming on Caribbean Coral Reefs

**DOI:** 10.1101/2021.05.12.443696

**Authors:** Colleen B. Bove, Laura Mudge, John F. Bruno

## Abstract

Anthropogenic climate change is rapidly altering the characteristics and dynamics of biological communities. This is especially apparent in marine systems as the world’s oceans are warming at an unprecedented rate, causing dramatic changes to coastal marine systems, especially on coral reefs of the Caribbean. We used three complementary ocean temperature databases (HadISST, Pathfinder, and OISST) to quantify change in thermal characteristics of Caribbean coral reefs over the last 150 years (1871–2020). These sea surface temperature (SST) databases included combined *in situ* and satellite-derived SST (HadISST, OISST), as well as satellite-only observations (Pathfinder) at multiple spatial resolutions. We also compiled a Caribbean coral reef database identifying 5,326 unique reefs across the region. We found that Caribbean reefs have warmed on average by 0.20 °C per decade since 1987, the calculated year that rapid warming began on Caribbean reefs. Further, geographic variation in warming rates ranged from 0.17 °C per decade on Bahamian reefs to 0.26 °C per decade on reefs within the Southern and Eastern Caribbean ecoregions. If this linear rate of warming continues, these already threatened ecosystems would warm by an *additional* 1.6 °C on average by 2100. We also found that marine heatwave (MHW) events are increasing in both frequency and duration across the Caribbean. Caribbean coral reefs now experience on average 5 MHW events annually, compared to 1 per year in the early 1980s. Combined, these changes have caused a dramatic shift in the composition and function of Caribbean coral reef ecosystems. If reefs continue to warm at this rate, we are likely to lose even the remnant Caribbean coral reef communities of today in the coming decades.

## Introduction

Greenhouse gas emissions are warming the planet, intensifying disturbances (e.g., fires and cyclonic storms), and modifying countless other aspects of the environment. This is causing extinctions, altering species composition, and degrading nearly every ecosystem on earth. Although we tend to think of surface warming as a terrestrial phenomenon, the oceans have stored about 93% of the additional retained heat since 1955 ^1,2^. The impacts of warming on marine communities are widespread, affecting a large range of taxa ^3^. Most marine species are ectothermic, so their body temperature matches that of the surrounding seawater. Therefore, warming increases their metabolism and subsequently their caloric demands, growth rates, behaviors, etc ^4,5^. This in turn has widespread effects on species interactions and the structure of marine food webs ^6^. Ocean heating has also been linked to disease outbreaks, the loss of foundation species that provide habitat structure (e.g., corals and kelps), reductions in primary production, and many other changes ^7^. The recent National Climate Assessment ^8^ described the well-documented effects of ocean warming as “ecosystem disruption” and concluded it will “intensify as ocean warming, acidification, deoxygenation, and other aspects of climate change increase.”

Warming episodes are driving unprecedented changes in coral reef ecosystem function and biodiversity globally ^9–11^. These changes on coral reefs are particularly evident on Caribbean reefs that have already experienced dramatic ecological shifts over the past several decades ^12,13^. Increasing SST on coral reefs has been associated with many negative ecological consequences, including coral bleaching ^14^, higher disease prevalence ^15^, increased mortality ^16^, and overall reductions in metabolic processes across marine taxa ^17,18^. While ocean warming is a global phenomenon, differences in the rate of warming can be highly localized ^19^, emphasizing the need to understand the thermal histories of the ecosystems most vulnerable to projected ocean warming.

The purpose of this study was to quantify the long-term (150 year) spatiotemporal trends in ocean temperature across the Greater Caribbean, and specifically on Caribbean coral reefs. We compiled three open-access sea surface temperature (SST) datasets (HadISST, Pathfinder SST, and OISST) for all mapped coral reef locations across the Greater Caribbean (**Figure 1**). Numerous previous studies have documented the anthropogenic heating of the ocean generally ^2,20^, and of coral reefs in particular ^19,21^. Our study builds on this work by focusing on coral reefs of the Greater Caribbean, updating the analysis through 2020, and by adding an assessment of coral reef marine heatwaves to the standard focus on spatiotemporal trends temperature.

**Figure 1.**
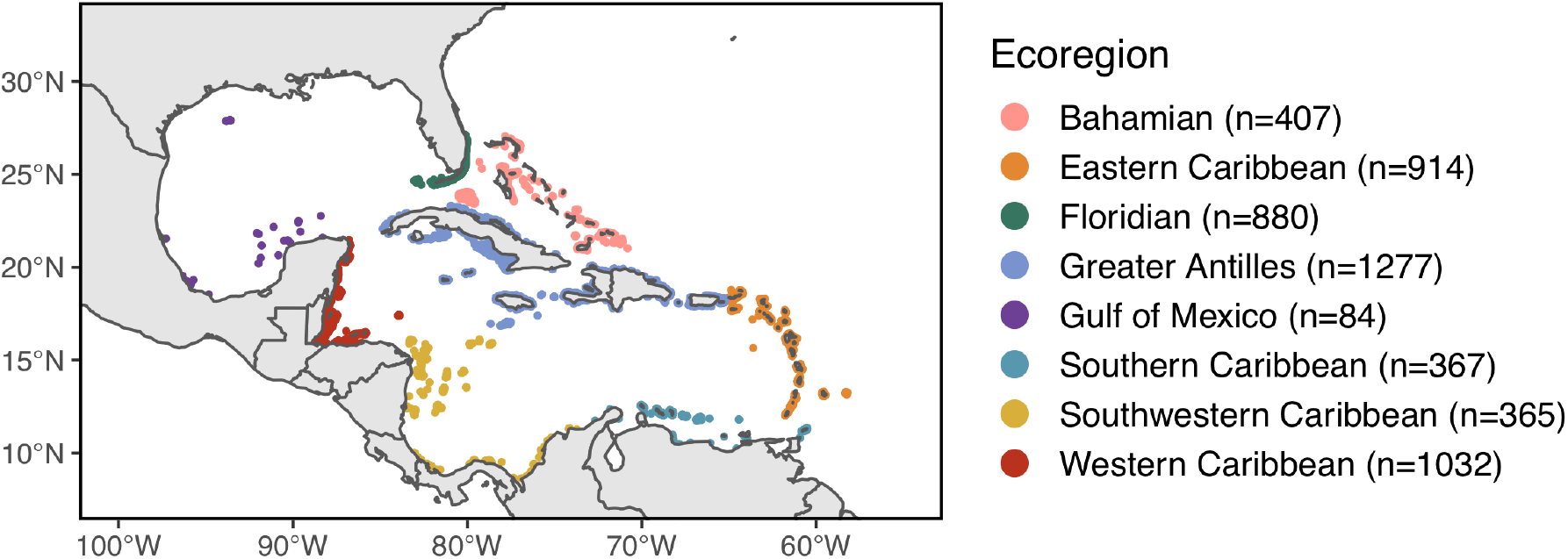
Caribbean coral reef site locations and ecoregion designation. The colour of each reef represents the designated ecoregion and n denotes the number of unique reef locations within that ecoregion.

## Results and Discussion

Our results indicate that Caribbean coral reefs continue to warm and that warming over the last 150 years was apparent every month (**Figure 2A**). In fact, recent spring temperatures often meet or exceed annual highs (typically in September) observed in the late 19th century. Caribbean reef surface temperatures were relatively stable in the mid 20th century (due to volcanic activity ^22^), then warming increased around 1987 (**Figures 2B; Supplementary Figure 1**). This timing is similar to long-term records of global ocean surface temperature and ocean heat content ^23^ (**Figure 2B**). We found that the average linear warming rate for the Caribbean reefs over the last 30 years (1987–2020; HadISST database) was 0.16 °C per decade (**Figure 2; Table 1**). Based on satellite data alone (**Supplementary Figures 2 and 3**; Pathfinder database), the average coral reef warming rate during this period was 0.20 °C per decade (**Table 1**). At this rate, the mean temperature on Caribbean reefs would be roughly 1.6 °C higher by 2100 assuming continued linear warming — and that is in addition to already realized warming.

**Figure 2.**
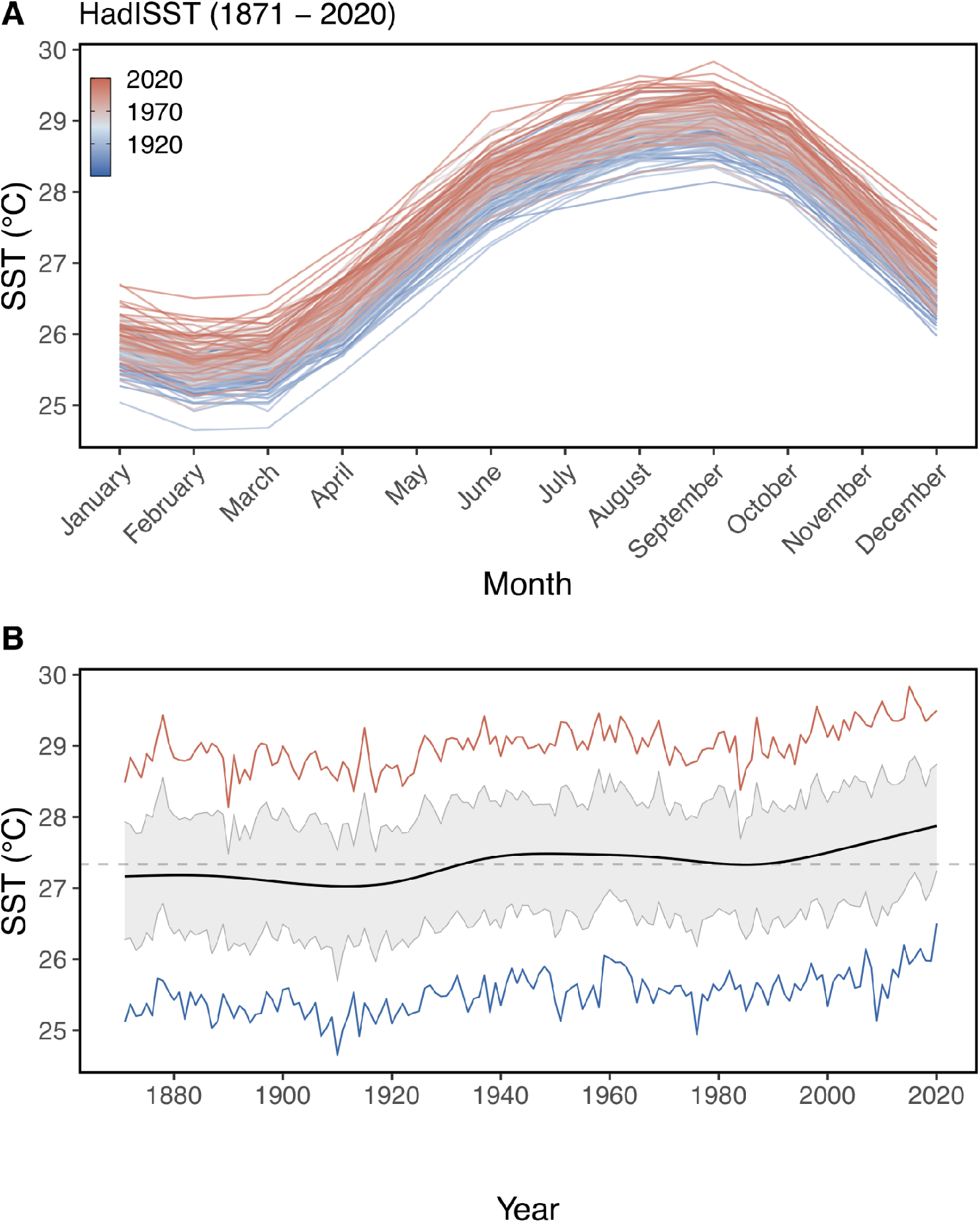
Historic SST trend on Caribbean coral reefs (1871–2020). Long-term SST records (HadISST) on Caribbean coral reefs depicting **A)** mean monthly SST each year (represented by line colour: blue to red) and **B)** GAM smoothed annual mean SST time (black line), annual maximum (red line), and annual minimum (blue line) SST. The grey dashed horizontal line denotes the overall mean SST for all sites over the entire period (27.3 °C) and the grey ribbon represents the 95% confidence interval around the annual mean SST through time.

**Table 1.**
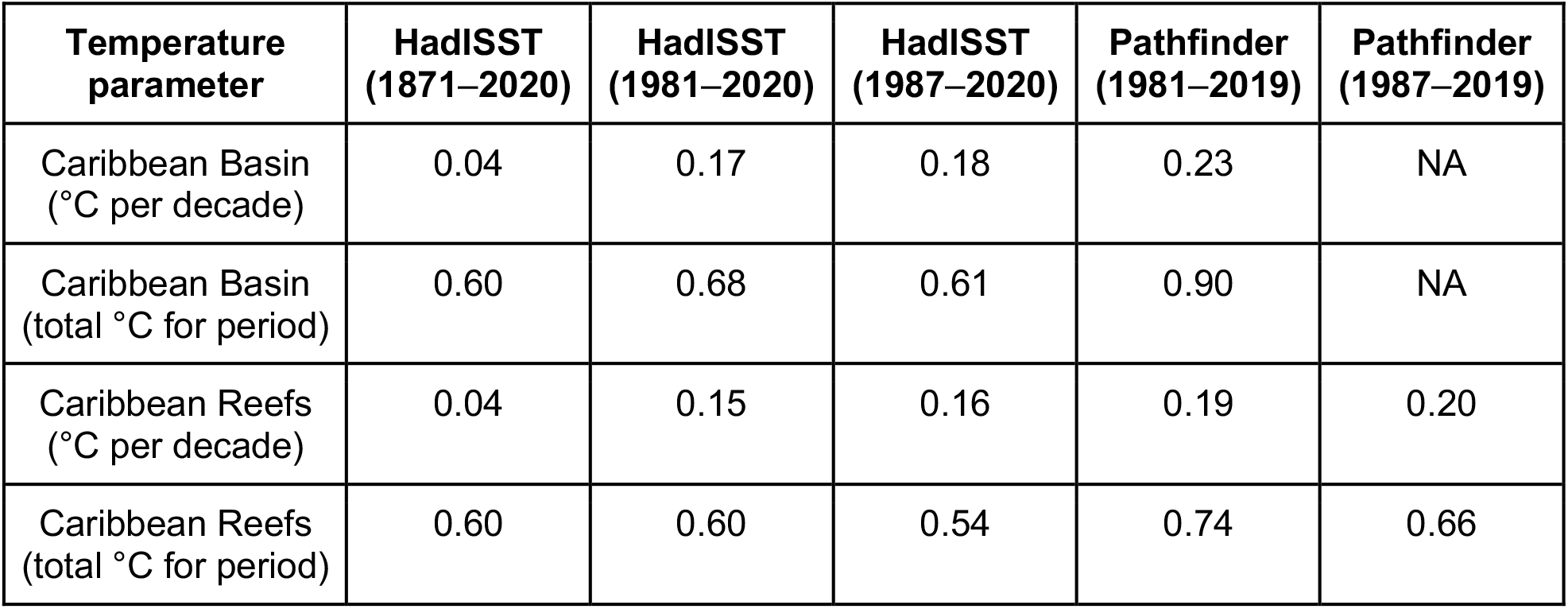
Estimated warming rates from both HadISST and Pathfinder databases for different temporal ranges. Values are means of all 5,326 reef locations included in the study. The year 1987 was estimated as the beginning of the most recent period of warming across all Caribbean coral reefs (**see Supplementary Figure 1**).

| Temperature parameter | HadISST (1871–2020) | HadISST (1981–2020) | HadISST (1987–2020) | Pathfinder (1981–2019) | Pathfinder (1987–2019) |
| --- | --- | --- | --- | --- | --- |
| Caribbean Basin (°C per decade) | 0.04 | 0.17 | 0.18 | 0.23 | NA |
| Caribbean Basin (total °C for period) | 0.60 | 0.68 | 0.61 | 0.90 | NA |
| Caribbean Reefs (°C per decade) | 0.04 | 0.15 | 0.16 | 0.19 | 0.20 |
| Caribbean Reefs (total °C for period) | 0.60 | 0.60 | 0.54 | 0.74 | 0.66 |

The observed warming rates of Caribbean coral reefs in this and previous studies report similar values, however, these rates are somewhat greater than estimates for the global ocean surface (**Table 2**). Winter et al. ^21^ reported a warming rate of about 0.25 °C per decade for the reef off La Parguera, southwestern Puerto Rico (1966–1995). Similarly, Hoegh-Guldberg ^24^ found the warming rate was 0.23 °C per decade (1981–1999) off the south coast of Jamaica. Kuffner et al. ^25^ described a remarkable temperature record collected by lighthouse keepers for five coral reefs off the Florida Keys starting in 1878 that also observed SST warming at 0.25 °C per decade between 1975 and 2006. It is reassuring (if surprising) that studies of vastly different scales and based on disparate methods, including *in situ* measurements (e.g., filling a bucket with seawater then measuring temperature by hand with a thermometer) ^21^, remote sensing via satellite ^19^, and databases based on both (this study) report similar warming rates. Moreover, Caribbean reef warming rates are approaching the CMIP5 RCP 8.5 model prediction of about 0.3 °C per decade ^26^. This “business as usual” emissions model is now viewed by many climate scientists as “unlikely”, especially for the end of this century. Yet, our results and previous work suggest that Caribbean coral reefs are already warming far more quickly than scenarios considered more likely (e.g., RCP 4.5).

**Table 2.**
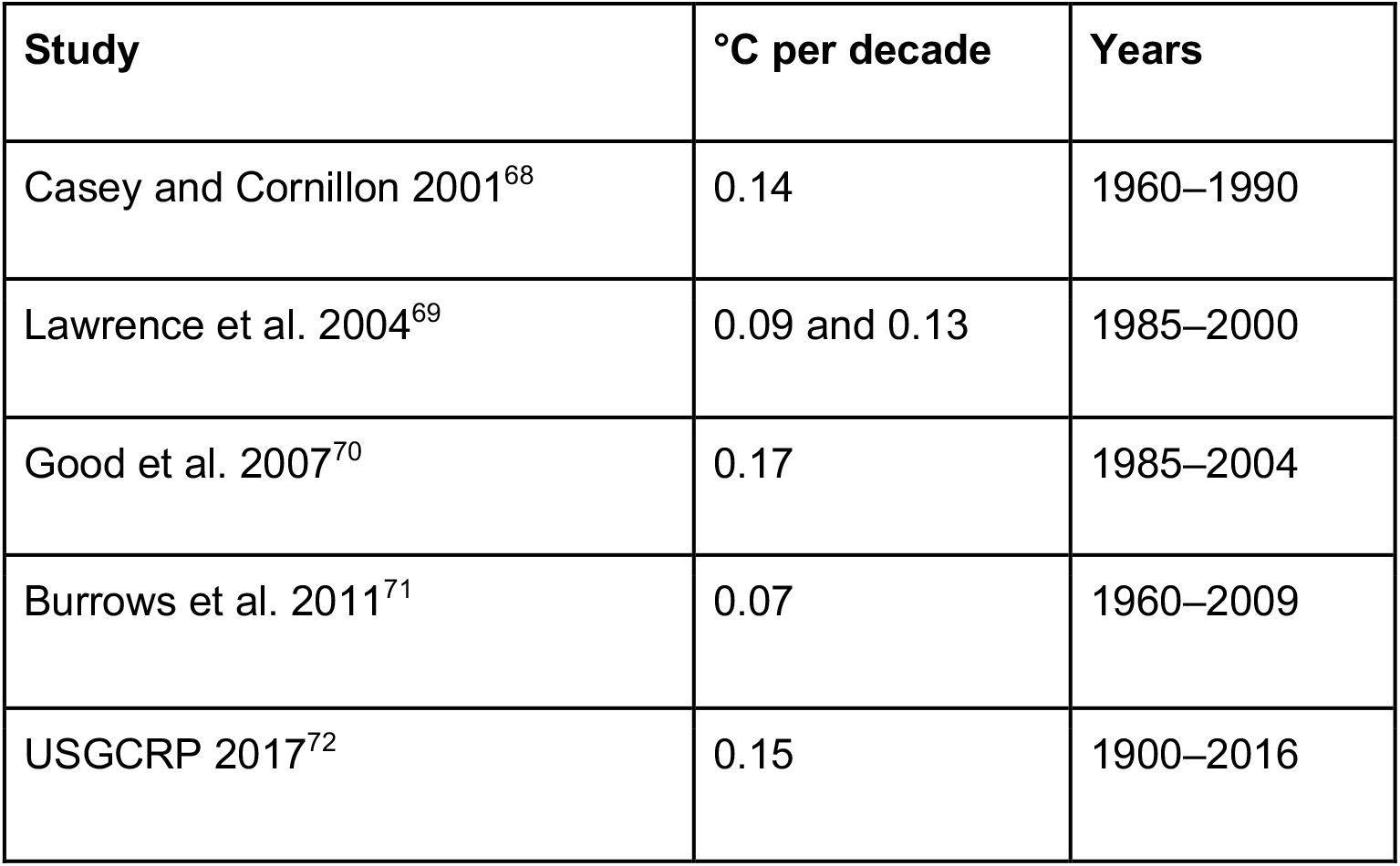
Published reports of global ocean surface warming rates.

| Study | °C per decade | Years |
| --- | --- | --- |
| Casey and Cornillon 2001<sup>68</sup> | 0.14 | 1960–1990 |
| Lawrence et al. 2004<sup>69</sup> | 0.09 and 0.13 | 1985–2000 |
| Good et al. 2007<sup>70</sup> | 0.17 | 1985–2004 |
| Burrows et al. 2011<sup>71</sup> | 0.07 | 1960–2009 |
| USGCRP 2017<sup>72</sup> | 0.15 | 1900–2016 |

We compared long-term coral reef temperature trends among eight previously-defined ecoregions within the Caribbean ^27^. Our results indicate that reefs within all ecoregions have clearly warmed since 1871 (**Figures 3**; **Supplementary Figure 4**), albeit at somewhat different rates. Our inflection point analysis determined that the initial year of rapid warming also varied among ecoregions. For example, rapid warming was identified in the Gulf of Mexico and Southern Caribbean starting in 1981, while rapid warming was not detected until 1999 in the Western Caribbean (**Figures 3; Supplementary Figure 4; Supplementary Table 2**). So, although the Western Caribbean began warming much later, recently it has warmed at a substantially higher rate of 0.24 °C per decade. Since recent rapid warming began in each region, warming rates ranged from about 0.17 °C per decade on Bahamian reefs to 0.26 °C per decade on reefs in the Southern and Eastern Caribbean (**Figure 3**; **Supplementary Table 2**). Overall, these warming rates translate to an increase in annual coral reef temperatures between 0.53 and 0.99 °C over the last 30 years.

**Figure 3.**
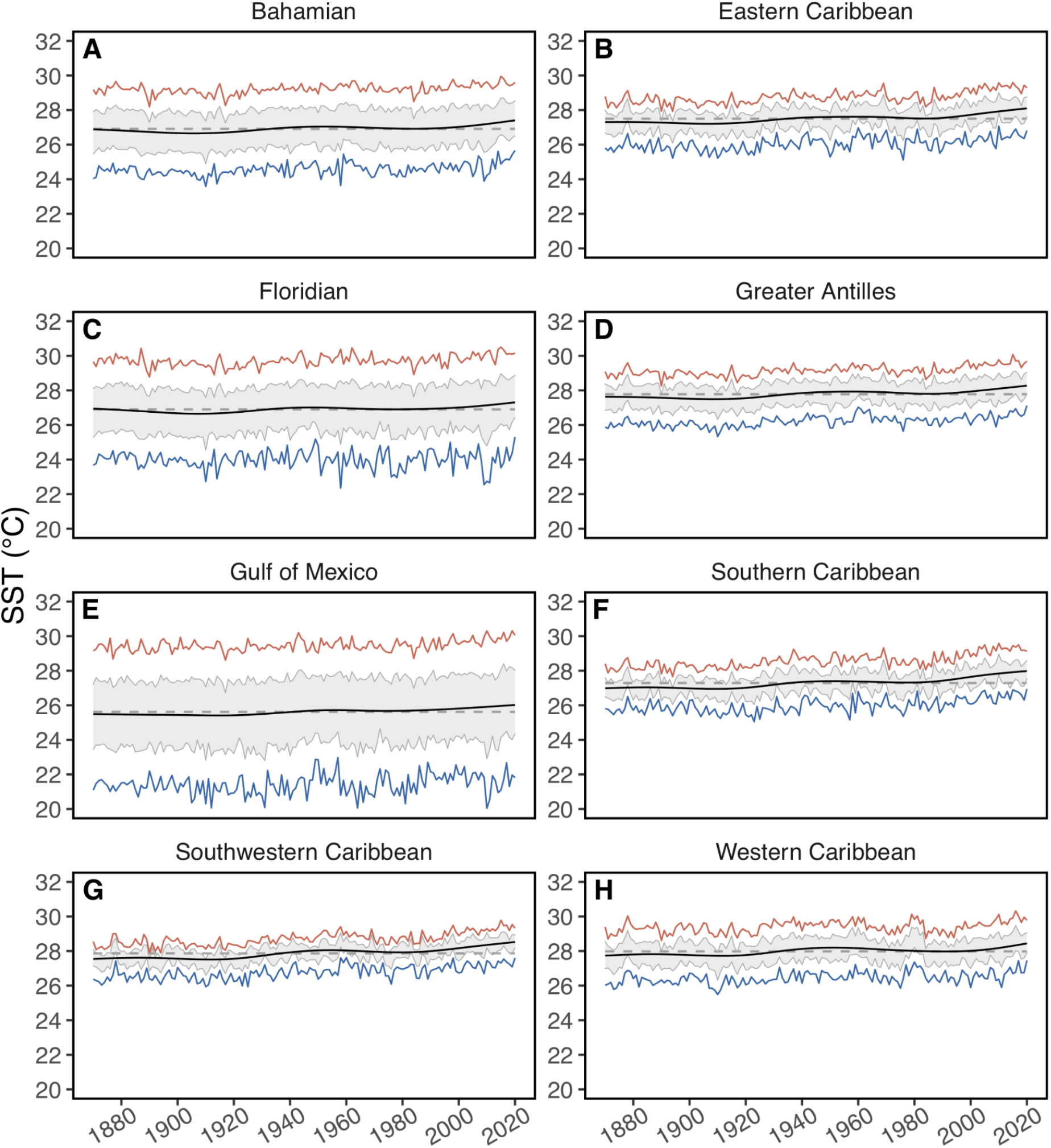
Historic SST trends on coral reefs within ecoregions (1871–2020). Long-term SST records (HadISST) on Caribbean coral reefs separated by ecoregion depicting GAM smoothed annual mean SST time (black line), annual maximum (red line), and annual minimum (blue line) SST. The grey dashed horizontal line denotes the mean SST over the entire period and the grey ribbon represents the 95% confidence interval around the true annual SST mean for each ecoregion.

Subregional patterns of warming across the Caribbean calculated here are similar to other SST parameters used for assessing thermal risk on coral reefs, such as degree heating weeks (accumulation of temperature anomalies exceeding the monthly maximum mean SST ^16^; DHW). Ecoregions within the Caribbean with faster rates of warming (e.g., Southern and Eastern Caribbean [mean 0.26 °C per decade], **Supplementary Table 2**) have also been reported to have some of the highest occurrences of weekly SST anomalies ^28^, maximum DHWs ^16^, and coral bleaching or mortality risk events ^29^. The variability in warming parameters across Caribbean ecoregions is likely a significant contributor to differences in coral cover among these regions since the mid 1990s ^30^.

While increasing SST on Caribbean coral reefs is clearly a major concern, warming is not limited to reef locations. Sea surface temperatures are rising across the entire Caribbean basin at a mean rate of 0.04 °C per decade since 1871 (HadISST data; **Supplementary Figure 6**) and more rapidly since 1981 at 0.23 °C per decade (Pathfinder SST data; **Figure 4A; Supplementary Figure 6**). This recent rate of ocean warming (0.23 °C per decade from 1981–2019) is similar to those previously calculated for the Caribbean basin. For example, Chollett et al. ^19^ reported a rate for the entire Caribbean and southeastern Gulf of Mexico of 0.27 °C per decade (1985–2009). It is clear that the Caribbean basin is experiencing rapid warming (**Figure 2B, 3, 4A; Supplementary Figures 6, and 7**). Additionally, rapid increases in ocean heat content across the region has been apparent ^31^, driving increases in other severe events such as increased storm activity and marine heatwaves (MHWs).

**Figure 4.**
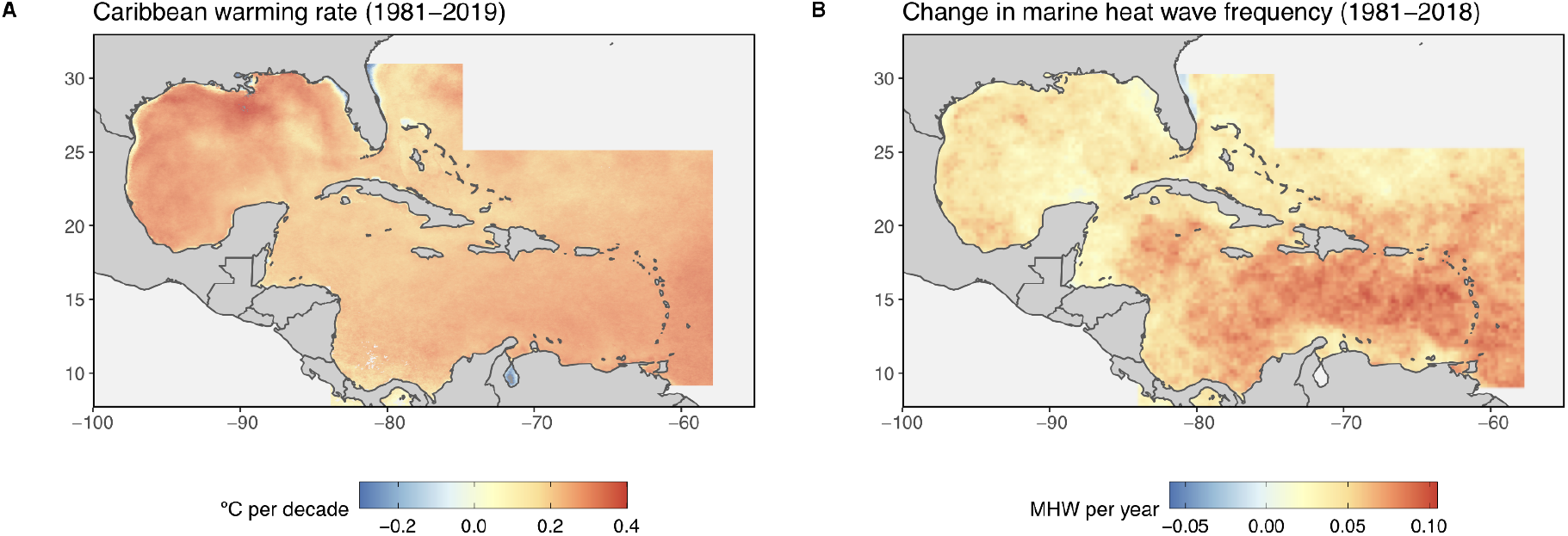
Warming patterns throughout the Caribbean Sea. Increasing warming events across the Caribbean depicted through **A)** rate of SST change (°C per decade) from 1981 to 2019 (Pathfinder; mean slope 0.23 ± 0.07 °C per decade) and **B)** increasing marine heatwave events (slope of counts per year). Grey ocean area was not included in these analyses.

We complemented our analyses of increasing Caribbean SST with an assessment of changes in marine heatwaves on Caribbean coral reefs. Marine heatwaves (MHW) are discrete warming events characterized by rate of onset, duration of event (five days or longer), and the intensity of warming ^32^. Over the past several decades, MHW have increased in both frequency and duration globally ^33^. Likewise, we found that heatwaves are occurring more frequently across the entire Caribbean basin (**Figure 4B; Supplementary Figure 5**) and on Caribbean coral reefs (**Figure 5A; Supplementary Tables 3 and 4**). The average frequency of MHW events on Caribbean coral reefs has increased from about 1 per year in the 1980s to almost 5 per year in the 2010s (**Figure 5A; Supplementary Table 4**), with current events lasting on average about 14 days each (**Figure 5B; Supplementary Table 4**). Additionally, the decadal mean return time (number of days elapsed since the last MHW event) has steadily declined from 377 days in the 1980s to just 111 days in the 2010s (**Figure 5C; Supplementary Table 4**), suggesting that Caribbean reef organisms and communities have approximately one third the time to recover from such thermal events.

**Figure 5.**
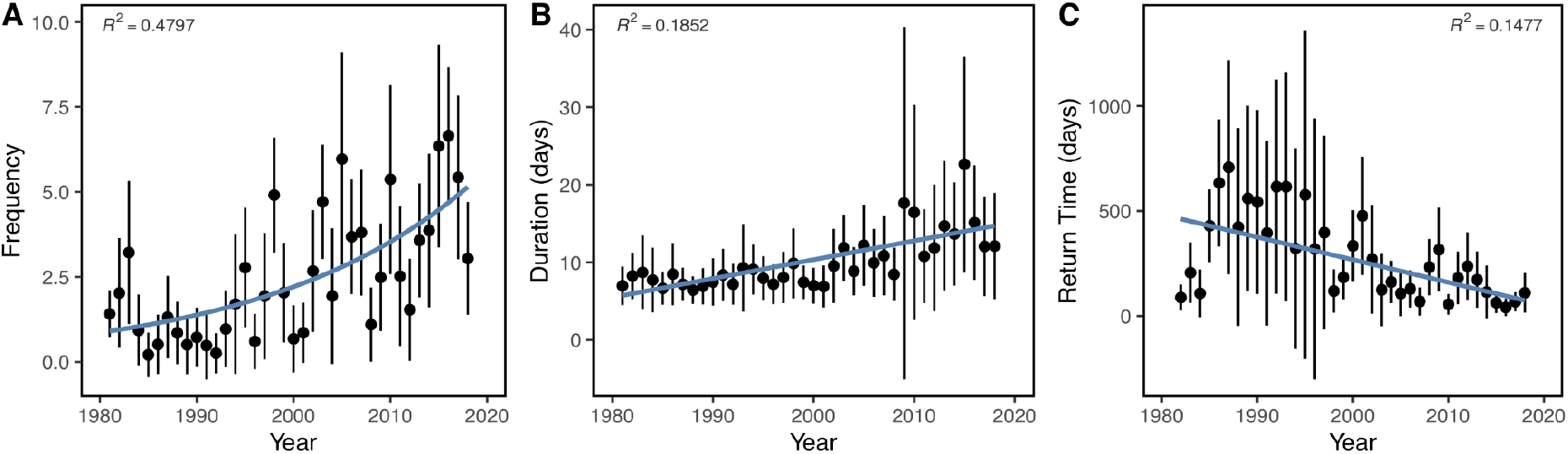
MHW trends (1981–2018) across Caribbean coral reefs. Temperature data are based on OISST gridded data to determine **A)** marine heat wave (MHW) frequency (number events per year) with Nagelkerke pseudo R^*2*^; **B)** MHW duration (number days per event) with linear model R^*2*^ ; and **C)** return time (number days per event) since the previous MHW event with linear model R^*2*^ reported. Points denote annual mean values (±SD) and blue lines represent linear (lm or glm) trends.

MHW trends are fairly consistent across Caribbean ecoregions (**Supplementary Figure 8**), however, coral reefs within the Eastern Caribbean are experiencing the greatest increase in MHW duration compared to the 1980s (**Supplementary Figure 8; Supplementary Table 4**), while the change in frequency of MHWs was the lowest on Western Caribbean reefs (**Supplementary Figure 8; Supplementary Table 4**). Such variations in MHW events across ecoregions further highlights differences in subregional warming patterns that impact the future success of coral reefs on both regional and local scales.

These acute MHW events, often lasting days to weeks, are believed to be the primary cause of mass-coral mortality across the Caribbean. In fact, a recent study demonstrated a positive correlation between MHW duration and the frequency of coral bleaching across the Caribbean and Gulf of Mexico ^34^. However, these severe heating events are often missed with current coral bleaching monitoring in place, such as DHWs, because of the cumulative nature of these alert systems ^35,36^. Even with longer MHW events being detected through these traditional coral bleaching monitoring systems, the ecological impact MHWs have on coral reef ecosystems are distinct. MHWs are known to trigger rapid bleaching and mass coral mortality on reefs that impact species with all levels of thermal tolerance and on highly localized spatial scales ^36,37^.

## Conclusions

The remarkable collapse of Caribbean coral reef ecosystems began when ocean heating in the region accelerated ∼30 years ago. Not only has warming been the primary driver of widespread coral mortality, but it has indirectly (via habitat loss) likely affected countless non-coral reef inhabitants when their own thermal tolerances were exceeded ^38–41^. And yet, this ecosystem transformation was caused by a mere 1 °C of warming (or substantially less in some ecoregions). This fact strongly suggests that we will lose what little remains in the coming years and decades if the anthropogenic heating of the Caribbean sea continues at this rate ^42^. Another 0.6 or 1.6 °C of warming (which based on our current warming rates would occur mid- and end-of-century, respectively) is likely a best case scenario at this point since oceans will continue to absorb excess heat even if global carbon neutrality goals are met. Despite the uncertainties of future ocean warming patterns, it is clear that the last 150 years of warming on Caribbean coral reefs have drastically altered these ecosystems and it will take significant global efforts to prevent the total loss of these remaining coral reefs.

## Methods

### Reef Locations

We compiled a Caribbean coral reef location database by sourcing latitude and longitude coordinates for known coral reef locations from the following sources: UNEP World Conservation Monitoring Center ^43^, the Global Coral Reef Monitoring Program ^44^, the Atlantic and Gulf Rapid Reef Assessment (AGRRA) ^45^, Reef Check (reefcheck.org), Florida’s Coral Reef Evaluation and Monitoring Program (CREMP) ^46^, the US Virgin Islands Territorial Coral Reef Monitoring Program (TCRMP) ^47^, and previously published survey data for the region ^30^. Sites were considered duplicates if they had the same GPS coordinates. In total, we identified 5,326 unique reefs across the Caribbean basin that were assigned to ecoregions based on the World Wildlife Fund (WWF) marine ecoregion classifications ^27^. Ecoregions contained between 84 (Gulf of Mexico) and 1,277 (Greater Antilles) reef locations (**Figure 1**). This database was used to assess SST and MWH trends across Caribbean coral reefs.

### Sea Surface Temperature Datasets

Our characterization of the thermal history of the Caribbean was based on three complementary ocean temperature datasets: HadISST from the United Kingdom Met Office ^48^, the Pathfinder satellite temperature records from NOAA/NASA ^49–51^, and NOAA’s daily Optimum Interpolation Sea Surface Temperature (OISST) ^52^. We combined the HadISST and Pathfinder databases to assess both long-term and high-resolution SST trends across the Caribbean and solely on coral reefs. Additionally, we used the OISST database to identify and assess marine heatwave (MHW) events.

We obtained monthly SST from 1871 to 2020 from the HadISST dataset ^53,54^ at a resolution of 1° grids across the Caribbean (0°N–40°N; 100°W–55°W). The HadISST data are based on a combination of temperature reconstruction and observational data to produce a long-term record of *in situ* measurements (typically from ships and buoys) and satellite-derived temperatures. HadISST uses *in situ* SST data from the Met Office Marine Data Bank (MDB) and is supplemented with data from the Comprehensive Ocean-Atmospheric Data Set (ICOADS) when missing from the MDB between the years 1871 and 1995. Satellite SST data from the Global Telecommunications System are included in the HadISST dataset after 1982, making this database ideal for long-term SST assessments ^54,55^..

Additionally, we calculated monthly mean SST at 4 km resolution from September 1981 to December 2019 using the twice daily (night and day) Pathfinder Version 5.3 database ^49,50^. Pathfinder SST data were clipped to the wider Caribbean constraints and quality filtered (quality four or greater ^50,51^) before the mean monthly SST value was calculated per grid cell and concatenated into a single netCDF file. The Pathfinder database is derived from measurements made by the Advanced Very High Resolution Radiometer (AVHRR) instruments aboard NOAA’s polar orbiting satellites that combine multiple passes of data. These data were provided by GHRSST and the NOAA National Centers for Environmental Information.

We used NOAA’s AVHRR 0.25° daily OISST data to determine sea surface temperature climatology and detect MHW events across the Caribbean basin from 1982 to 2018 ^52,56^. OISST data are constructed by combining both *in situ* (collected via ships and buoys) and infrared satellite (AVHRR instruments) SST, applying a bias adjustment to satellite and ship observation data, and filling gaps as necessary through interpolation. The combination of high resolution and multiple sources of SST observations is ideal for identifying MHWs, and is frequently used for such assessments ^32,57^.

### Historic SST Assessment

We examined the rate of SST change through time for the wider Caribbean region using the HadISST (1871–2020) and Pathfinder (1981–2019) databases (described above) to evaluate historic SST trends. A simple linear model was applied to each grid cell through time (months each year) of both datasets to calculate the slope and significance of SST increases over time for the full region. Additionally, we extracted SST from the HadISST and Pathfinder datasets using a compiled Caribbean coral reef location database (*raster* package; version 2.9-23 ^58^). The use of satellite-derived SST measurements to represent benthic temperature patterns has been frequently assessed via comparisons of different databases with *in situ* logger measurements at the same locations. While these studies report that satellite-derived measurements often underestimate *in situ* temperatures ^59,60^, the consensus is that satellite databases accurately reflect temperature conditions on many coral reefs at depth ^60–62^ and are a useful tool for identifying global warming signals ^53^. Monthly SST measurements were averaged across all reef locations to assess historic SST trends on Caribbean coral reefs per sampling period (month) as well as within individual ecoregions. We assessed annual SST on coral reefs since 1871 using a generalized additive model (GAM) with a cubic regression spline for year and a cyclic cubic regression spline for month to smooth temperature variability through time for better assessment of temporal SST trends on reefs. We then identified the year in which annual warming rates significantly increased based on the first derivative of the GAM curve for the entire region as well as each ecoregion ^63^. These years were used to calculate the mean warming rate since the identified inflection points for coral reefs within the eight ecoregions. All analyses and visualizations were conducted using R version 3.6.1 ^64^.

### Marine Heatwaves Classification

Marine heatwaves (MHW) are defined as anomalously warm sea-surface temperature events, with temperatures warmer than historical climatology for that specific time and location, lasting for at least a 5 day period ^32^. We used NOAA’s OISST data (described above) to identify and assess MHWs across the wider Caribbean from 1982–2018 ^65^. If successive events had less than two days between them, they were considered to be the same event. We used the *heatwaveR* package in R (version 0.4.4) ^66^ and code provided from the marine heatwave working group (www.marineheatwaves.org) to identify marine heatwave events from our OISST dataset and calculate heatwave metrics (**Supplementary Table 1**).

For each pixel, we aggregated a list of distinct MHW events between 1982 and 2018, which included the start and end date of the event (see **Supplementary Table 1** for all MHW properties obtained). We calculated frequency of events per pixel by summing the total number of MHW events per year and the total number of MHW days as the sum of the duration of all events that year. Return time was calculated as the number of days elapsed between each unique MHW event. All other metrics were averaged annually.

### Caribbean Marine Heatwave Trends

To evaluate MHW trends across the Caribbean basin, we created an annual time series for each MHW pixel that contains the number of distinct MHW events, total number of MHW days experienced, the average maximum (peak) intensity (°C) for all events that year, the average duration (days) of all events that year, the average onset rate for all events that year (°C per day), and the average time elapsed since a previous event (days). We then calculated a history of MHW events for each coral reef location using a nearest neighbor analysis to match reef locations to MHW events identified from the OISST grids.

We quantified the temporal trends for each MHW metric using ordinary least squares (OLS) models, except for event frequency and total number of MHW days, as these variables are count data and best analyzed using generalized linear models (GLM) (*lme4*; version 1.1-23) ^67^. The response variables for OLS models were log transformed to meet model assumptions when necessary. For MHW trends on coral reefs, the level of observation was each unique MHW pixel that overlapped coral reef habitat and we modeled trends for each ecoregion to account for potential spatial differences in MHW metrics across the Caribbean basin. Significance and R^2^ values are obtained to evaluate the strength of each trend. For GLM trends, we report the Nagelkerke pseudo R^2^ value.

## Acknowledgements

This paper was funded in part by a grant from the National Science Foundation to JFB (OCE 1737071). We thank the following data providers: Atlantic Gulf Rapid Reef Assessment (AGRRA) contributors and data managers; the Reef Check Foundation, UNEP World Conservation Monitoring Center, the Global Coral Reef Monitoring Program, Florida’s Coral Reef Evaluation and Monitoring Program (CREMP), and the US Virgin Islands Territorial Coral Reef Monitoring Program (TCRMP). We thank Matthew Kendall for his assistance with cluster development and support with acquiring satellite data and S. Williams, H. Reich, J. Rippe, and J. Baumann for their helpful feedback on early drafts of the manuscript.

## Data Accessibility

HadISST data can be accessed at www.metoffice.gov.uk/hadobs/hadisst/, Pathfinder data can be accessed at www.ncei.noaa.gov/data/oceans/pathfinder/Version5.3/L3C/, and OISST data can be accessed at www.ncdc.noaa.gov/oisst/data-access. All data and code compiled for this manuscript can be freely accessed on GitHub (github.com/seabove7/CaribbeanSST) and Zendo (DOI: 10.5281/zenodo.4751658), including links to the compiled databases used here from the three sources listed above.

## Author Contributions

All authors conceived the idea and contributed intellectually to its development. CBB led the SST analyses and LM led the MHW analyses. CBB and JFB led the manuscript preparation with contributions from LM.

## Literature Cited

1. Cheng, L. et al. Improved estimates of ocean heat content from 1960 to 2015. Sci. Adv. 3, e1601545 (2017).

2. Levitus, S. et al. World ocean heat content and thermosteric sea level change (0-2000 m), 1955-2010: WORLD OCEAN HEAT CONTENT. Geophys. Res. Lett. 39, /a-n/a (2012).

3. Poloczanska, E. S. et al. Global imprint of climate change on marine life. Nature Climate Change 3, 919–925 (2013).

4. Seibel, B. A. & Drazen, J. C. The rate of metabolism in marine animals: environmental constraints, ecological demands and energetic opportunities. Philosophical Transactions of the Royal Society B: Biological Sciences 362, 2061–2078 (2007).

5. Newell, R. C. & Branch, G. M. The Influence of Temperature on the Maintenance of Metabolic Energy Balance in Marine Invertebrates. in Advances in Marine Biology (eds. Blaxter, J. H. S., Russell, F.S. & Yonge, M.) vol. 17 329–396 (Academic Press, 1980).

6. Bruno, J. F., Carr, L. A. & O’Connor, M. I. Exploring the role of temperature in the ocean through metabolic scaling. Ecology 96, 3126–3140 (2015).

7. Harvell, C. D. et al. Climate warming and disease risks for terrestrial and marine biota. Science 296, 2158–2162 (2002).

8. Pershing, A. et al. Oceans and marineresources: Impacts, risks, and adaptation in the United States. The fourth National Climate Assessment, Volume II, chapter 9. https://nca2018.globalchange.gov/chapter/9/ (2018) xdoi:10.7930/NCA4.2018.CH9.

9. Robinson, J. P. W., Wilson, S. K., Jennings, S. & Graham, N. A. J. Thermal stress induces persistently altered coral reef fish assemblages. Global Change Biology 25, 2739–2750 (2019).

10. Hughes, T. P. et al. Global warming transforms coral reef assemblages. Nature 556, 492–496 (2018).

11. Graham, N. A. J., Jennings, S., MacNeil, M. A., Mouillot, D. & Wilson, S. K. Predicting climate-driven regime shifts versus rebound potential in coral reefs. Nature 518, 94–97 (2015).

12. Gardner, T. A., Côté, I. M., Gill, J. A., Grant, A. & Watkinson, A. R. Long-Term Region-Wide Declines in Caribbean Corals. Science 301, 958–960 (2003).

13. Alvarez-Filip, L., Dulvy, N. K., Gill, J. A., Côté, I.M. & Watkinson, A. R. Flattening of Caribbean coral reefs: region-wide declines in architectural complexity. Proceedings of the Royal Society B: Biological Sciences 276, 3019–3025 (2009).

14. Hoegh-Guldberg, O. Climate change, coral bleaching and the future of the world’s coral reefs. Mar. Freshwater Res. (1999) doi:10.1071/MF99078.

15. Woesik, R. van & Randall, C. J. Coral disease hotspots in the Caribbean. Ecosphere 8, e01814 (2017).

16. Eakin, C. M. et al. Caribbean Corals in Crisis: Record Thermal Stress, Bleaching, and Mortality in 2005. PLoS One 5, (2010).

17. Bahr, K. D., Rodgers, K. S. & Jokiel, P. L. Ocean warming drives decline in coral metabolism while acidification highlights species-specific responses. Marine Biology Research 14, 924–935 (2018).

18. Harianto, J., Nguyen, H. D., Holmes, S. P. & Byrne, M. The effect of warming on mortality, metabolic rate, heat-shock protein response and gonad growth in thermally acclimated sea urchins (Heliocidaris erythrogramma). Mar Biol 165, 96 (2018).

19. Chollett, I., Müller-Karger, F. E., Heron, S. F., Skirving, W. & Mumby, P. J. Seasonal and spatial heterogeneity of recent sea surface temperature trends in the Caribbean Sea and southeast Gulf of Mexico. Marine Pollution Bulletin 64, 956–965 (2012).

20. Cheng, L., Abraham, J., Hausfather, Z. & Trenberth, K. E. How fast are the oceans warming? Science 363, 128–129 (2019).

21. Winter, A., Appledoorn, R. S., Bruckner, A., Williams, E.H. Jr. & Goenaga, C. Sea surface temperatures and coral reef bleaching off La Parguera, Puerto Rico (northeastern Caribbean Sea). Coral Reefs 17, 377–382 (1998).

22. Gill, J. A., Watkinson, A. R., McWilliams, J. P. & Cote, I. M. Opposing forces of aerosol cooling and El Nino drive coral bleaching on Caribbean reefs. Proceedings of the National Academy of Sciences 103, 18870–18873 (2006).

23. Barton, A. D. & Casey, K. S. Climatological context for large-scale coral bleaching. Coral Reefs 24, 536–554 (2005).

24. Hoegh-Guldberg, O. Climate change, coral bleaching and the future of the world’s coral reefs. Marine and Freshwater Research 50, 839–866 (1999).

25. Kuffner, I. B., Lidz, B. H., Hudson, J. H. & Anderson, J. S. A Century of Ocean Warming on Florida Keys Coral Reefs: Historic In Situ Observations. Estuaries and Coasts 38, 1085–1096 (2015).

26. Bruno, J. F. et al. Climate change threatens the world’s marine protected areas. Nature Clim Change 8, 499–503 (2018).

27. Spalding, M. D. Marine Ecoregions of the World: a bioregionalization of coast and shelf areas: Working notes, 2010. (2010).

28. Selig, E. R., Casey, K. S. & Bruno, J. F. New insights into global patterns of ocean temperature anomalies: implications for coral reef health and management. Global Ecology and Biogeography 19, 397–411 (2010).

29. Muñiz-Castillo, A. I. et al. Three decades of heat stress exposure in Caribbean coral reefs: a new regional delineation to enhance conservation. Sci Rep 9, (2019).

30. Schutte, V., Selig, E. & Bruno, J. Regional spatio-temporal trends in Caribbean coral reef benthic communities. Mar. Ecol. Prog. Ser. 402, 115–122 (2010).

31. Cheng, L. et al. Upper Ocean Temperatures Hit Record High in 2020. Adv. Atmos. Sci. (2021) doi:10.1007/s00376-021-0447-x.

32. Hobday, A. J. et al. A hierarchical approach to defining marine heatwaves. Progress in Oceanography 141, 227–238 (2016).

33. Oliver, E. C. J. et al. Longer and more frequent marine heatwaves over the past century. Nature Communications 9, 1324 (2018).

34. Smale, D. A. et al. Marine heatwaves threaten global biodiversity and the provision of ecosystem services. Nature Climate Change 9, 306–312 (2019).

35. Liu, G. et al. NOAA Coral Reef Watch’s 5km Satellite Coral Bleaching Heat Stress Monitoring Product Suite Version 3 and Four-Month Outlook Version 4. 32, 7 (2017).

36. Fordyce, A. J., Ainsworth, T. D., Heron, S. F. & Leggat, W. Marine Heatwave Hotspots in Coral Reef Environments: Physical Drivers, Ecophysiological Outcomes, and Impact Upon Structural Complexity. Front. Mar. Sci. 6, (2019).

37. Le Nohaïc, M. et al. Marine heatwave causes unprecedented regional mass bleaching of thermally resistant corals in northwestern Australia. Scientific Reports 7, 14999 (2017).

38. Habary, A., Johansen, J. L., Nay, T. J., Steffensen, J. F. & Rummer, J. L. Adapt, move or die – how will tropical coral reef fishes cope with ocean warming? Global Change Biology 23, 566–577 (2017).

39. Stuart-Smith, R. D., Brown, C. J., Ceccarelli, D. M. & Edgar, G. J. Ecosystem restructuring along the Great Barrier Reef following mass coral bleaching. Nature 560, 92–96 (2018).

40. Jones, G. P., McCormick, M. I., Srinivasan, M. & Eagle, J. V. Coral decline threatens fish biodiversity in marine reserves. Proc Natl Acad Sci USA 101, 8251–8253 (2004).

41. Pratchett, M. S. et al. Effects of climate-induced coral bleaching on coral-reef fishes — ecological and economic consequences. in Oceanography and marine biology: an annual review vol. 46 251–296 (CRC Press, 2008).

42. Frieler, K. et al. Limiting global warming to 2 °C is unlikely to save most coral reefs. Nature Climate Change 3, 165–170 (2013).

43. UNEP-WCMC, WorldFish Centre, WRI & TNC. Global distribution of warm-water coral reefs, compiled from multiple sources including the Millennium Coral Reef Mapping Project. UN Environment World Conservation Monitoring Centre Version 4.0, (2018).

44. Jackson, E. J., Donovan, M., Cramer, K. & Lam, V. Status and trends of Caribbean coral reefs: 1970-2012. 306 (2012).

45. Marks, K. W. AGRRA Database, version (2018-03). (2018).

46. Porter, J. & Stossel, M. Coral Reef Evaluation and Monitoring Project Florida Keys 2001. (2020).

47. Smith, T. et al. The United States Virgin Islands Territorial Coral Reef Monitoring Program. 2016 Annual Report. (2016).

48. Rayner, N. A. Global analyses of sea surface temperature, sea ice, and night marine air temperature since the late nineteenth century. J. Geophys. Res. 108, 4407 (2003).

49. Korak, S. et al. AVHRR Pathfinder version 5.3 level 3 collated (L3C) global 4km sea surface temperature for 1981-Present. (2018) doi:10.7289/V52J68XX.

50. Casey, K. S., Brandon, T. B., Cornillon, P. & Evans, R. The Past, Present, and Future of the AVHRR Pathfinder SST Program. in Oceanography from Space (eds. Barale, V., Gower, J.F.R. & Alberotanza, L.) 273–287 (Springer Netherlands, 2010). doi:10.1007/978-90-481-8681-5_16.

51. Kilpatrick, K. A., Podestá, G.P. & Evans, R. Overview of the NOAA/NASA advanced very high resolution radiometer Pathfinder algorithm for sea surface temperature and associated matchup database. Journal of Geophysical Research: Oceans 106, 9179– 9197 (2001).

52. Reynolds, R. W. et al. Daily High-Resolution-Blended Analyses for Sea Surface Temperature. Journal of Climate 20, 5473–5496 (2007).

53. Kent, E. C. & Kennedy, J. J. Historical Estimates of Surface Marine Temperatures. Annual Review of Marine Science 13, 283–311 (2021).

54. Sheppard, C. & Rayner, N. A. Utility of the Hadley Centre sea ice and sea surface temperature data set (HadISST1) in two widely contrasting coral reef areas. Marine Pollution Bulletin 44, 303–308 (2002).

55. Lough, J. M., Anderson, K. D. & Hughes, T. P. Increasing thermal stress for tropical coral reefs: 1871–2017. Scientific Reports 8, 6079 (2018).

56. Reynolds, R. W., Rayner, N. A., Smith, T. M., Stokes, D. C. & Wang, W. An Improved In Situ and Satellite SST Analysis for Climate. Journal of Climate 15, 1609–1625 (2002).

57. Couch, C. S. et al. Mass coral bleaching due to unprecedented marine heatwave in Papahānaumokuākea Marine National Monument (Northwestern Hawaiian Islands). PLOS ONE 12, e0185121 (2017).

58. Hijmans, R. J. raster: Geographic Data Analysis and Modeling. (2019).

59. Castillo, K. D. & Lima, F. P. Comparison of in situ and satellite-derived (MODIS-Aqua/Terra) methods for assessing temperatures on coral reefs. Limnology and Oceanography: Methods 8, 107–117 (2010).

60. Gomez, A. M. et al. Comparison of Satellite-Based Sea Surface Temperature to In Situ Observations Surrounding Coral Reefs in La Parguera, Puerto Rico. Journal of Marine Science and Engineering 8, 453 (2020).

61. Claar, D. C., Cobb, K. M. & Baum, J. K. In situ and remotely sensed temperature comparisons on a Central Pacific atoll. Coral Reefs 38, 1343–1349 (2019).

62. Bruno, J. F., Siddon, C. E., Witman, J. D., Colin, P. L. & Toscano, M. A. El Niño related coral bleaching in Palau, Western Caroline Islands. Coral Reefs 20, 127–136 (2001).

63. Bennion, H., Simpson, G. L. & Goldsmith, B. J. Assessing degradation and recovery pathways in lakes impacted by eutrophication using the sediment record. Front. Ecol. Evol. 3, (2015).

64. R Core Team. R: A Language and Environment for Statistical Computing. (R Foundation for Statistical Computing, 2018).

65. Banzon, V., Smith, T. M., Steele, M., Huang, B. & Zhang, H.-M. Improved Estimation of Proxy Sea Surface Temperature in the Arctic. J. Atmos. Oceanic Technol. 37, 341–349 (2020).

66. Schlegel, R. W. & Smit, A. J. heatwaveR: A central algorithm for the detection of heatwaves and cold-spells. Journal of Open Source Software 3, 821 (2018).

67. Bates, D. & Maechler, M. Fitting Linear Mixed-Effects Models Using {lme4}. Journal of Statistical Software 67, 1–48 (2015).

68. Casey, K. S. & Cornillon, P. Global and regional sea surface temperature trends. Journal of Climate 14, 3801–3818 (2001).

69. Lawrence, S. P., Llewellyn-Jones, D. T. & Smith, S. J. The measurement of climate change using data from the Advanced Very High Resolution and Along Track Scanning Radiometers. J. Geophys. Res. 109, /a-n/a (2004).

70. Good, S. A., Corlett, G. K., Remedios, J. J., Noyes, E. J. & Llewellyn-Jones, D. T. The global trend in sea surface temperature from 20 years of Advanced Very High Resolution Radiometer Data. Journal of Climate 20, 1255–1264 (2007).

71. Burrows, M. T. et al. The Pace of Shifting Climate in Marine and Terrestrial Ecosystems. Science 334, 652–655 (2011).

72. USGCRP. Climate Science Special Report. 1–470 https://science2017.globalchange.gov/chapter/executive-summary/.

