## Supplemental Figures and Tables for "One Hundred and Fifty Years of Warming on Caribbean Coral Reefs"

**Supplemental Materials for manuscript:** *One Hundred and Fifty Years of Warming on Caribbean Coral Reefs*

Colleen B. Bove[1, 2*], Laura Mudge[1,3], and John F. Bruno[1]
[1]The Department of Biology, The University of North Carolina at Chapel Hill, Chapel Hill, North Carolina, 27599-3280 USA
[2]The Department of Biology, Boston University, Boston, Massachusetts, 02215 USA
[3]Integral Consulting Inc., Annapolis, Maryland 21401 USA

### Supplemental Figures

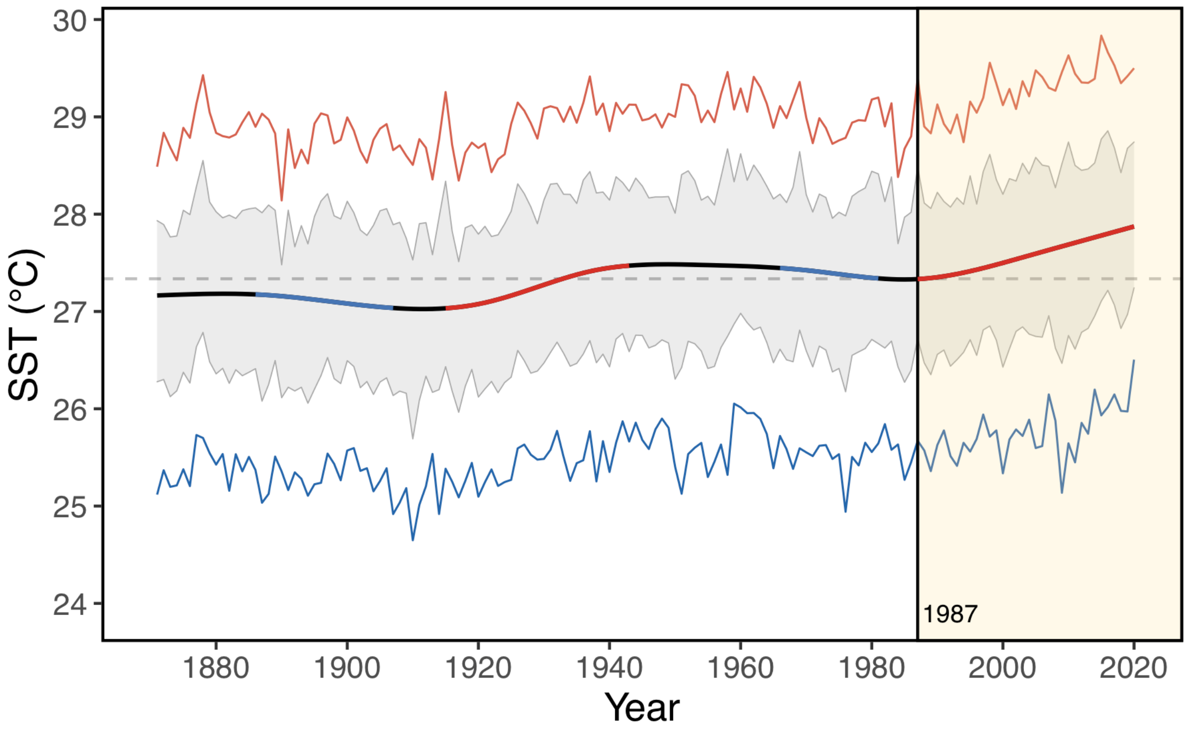


**Supplemental Figure 1.** Historic SST trend on Caribbean coral reefs (1871–2020). Long-term SST records (HadISST) on Caribbean coral reefs depicting significant changes in warming rate based on the first derivative of the GAM slope. The GAM smoothed annual mean SST time is represented by the black line with significantly positive (red) or negative (blue) slopes identified over the curve. The most recent significant warming event began in 1987 and is highlighted in the yellow box. The annual maximum (red line) and annual minimum (blue line) SST are also depicted, along with the overall mean SST for all sites over the entire period (27.3 °C; grey dashed line). The grey ribbon represents the 95% confidence interval around the true annual SST mean through time.

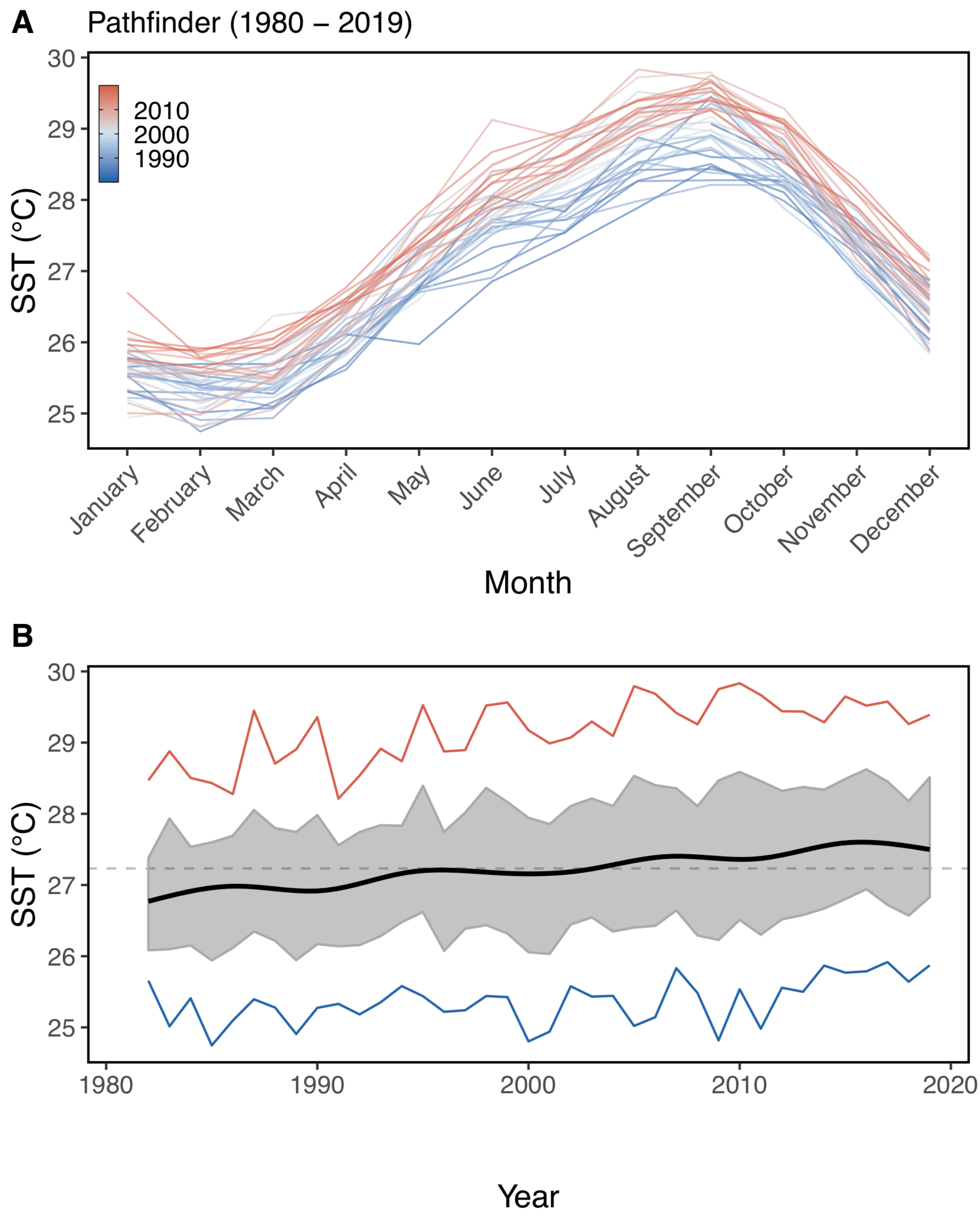


**Supplemental Figure 2.** Historic SST records (1981–2019; Pathfinder) on Caribbean coral reefs depicting **A)** mean monthly SST each year (represented by line colour: blue to red) and **B)** GAM smoothed annual mean SST time (black line), annual maximum (red line), and annual minimum (blue line) SST. The grey dashed horizontal line denotes the overall mean SST for all sites over the entire period (27.23 °C) and the grey ribbon represents the 95% confidence interval around the true annual SST mean through time.

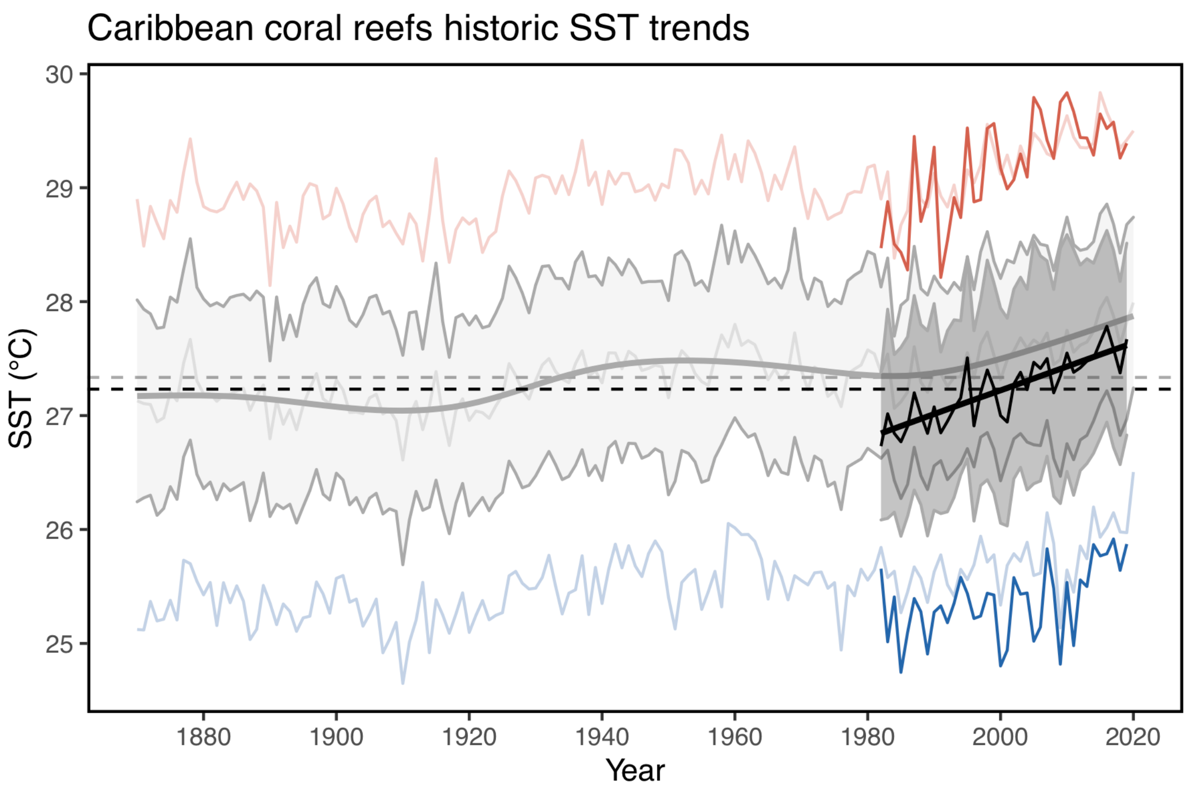


**Supplemental Figure 3.** Comparison of HadISST (1871–2020) and Pathfinder (1981–2019) SST recorded on Caribbean coral reef locations. The high-resolution Pathfinder is represented as darker data over the long-term HadISST. Both datasets are represented by GAM smoothed annual mean SST time (solid line), annual maximum (red line), and annual minimum (blue line) SST. The dashed horizontal line denotes the overall mean SST for all sites over the entire period and the grey ribbon represents the 95% confidence interval around the true annual SST mean through time.

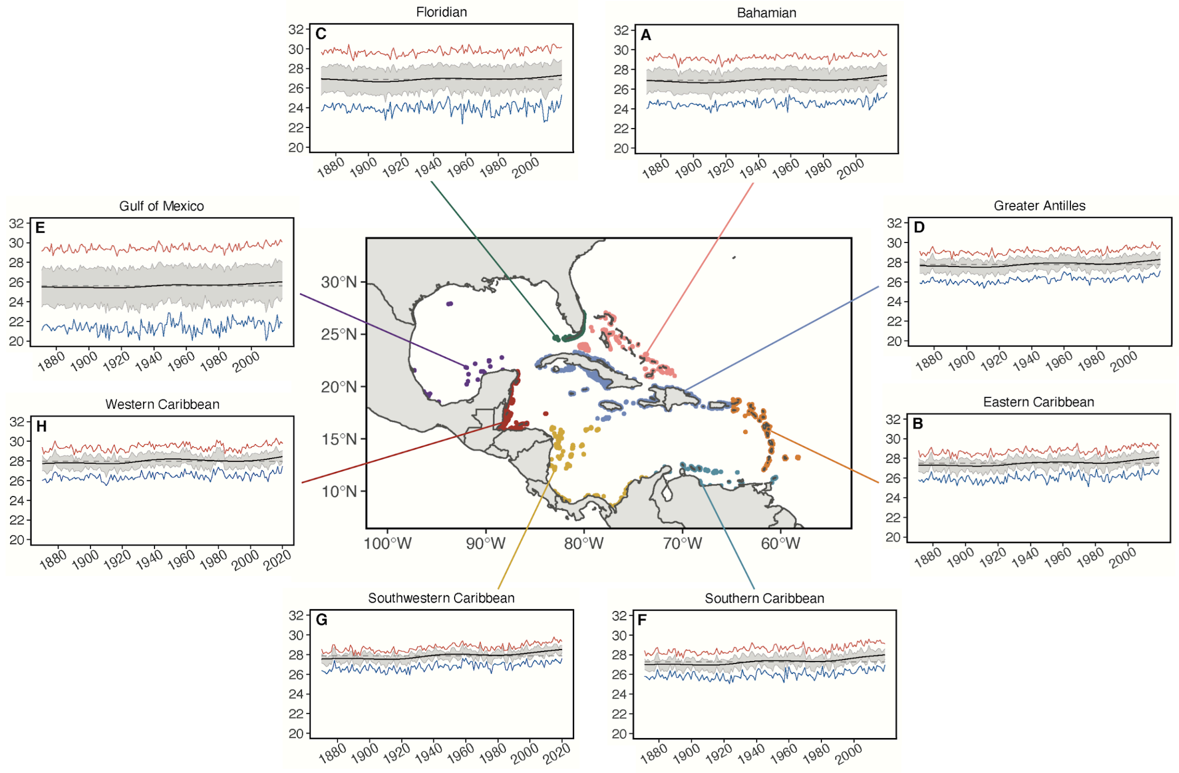


**Supplemental Figure 4**: Historic SST trends on coral reefs within Caribbean ecoregions (1871-2020) with corresponding reef locations (see **Figures 1, 3** in the main text). The colour of each reef location and box around long-term SST (HadISST) plots represent the designated ecoregion. Plots depict SST data with GAM smoothed annual mean SST time (black line), annual maximum (red line), and annual minimum (blue line) SST. The grey dashed horizontal line denotes the mean SST over the entire period and the grey ribbon represents the 95% confidence interval around the true annual SST mean for the **A)** Bahamian, **B)** Eastern Caribbean, **C)** Floridian, **D)** Greater Antilles, **E)** Gulf of Mexico, **F)** Southern Caribbean, **G)** Southwestern Caribbean, and **H)** Western Caribbean ecoregions.

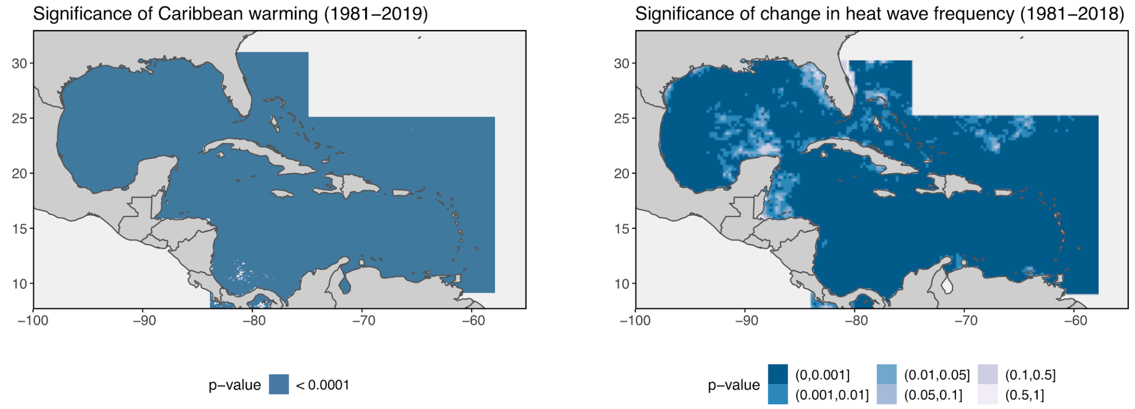


**Supplemental Figure 5**: Significance of **A)** rate of SST change (°C per decade) and **B)** number of marine heatwave events per year across the Caribbean depicted in **Figure 4**. Grey ocean area was not included in these analyses.

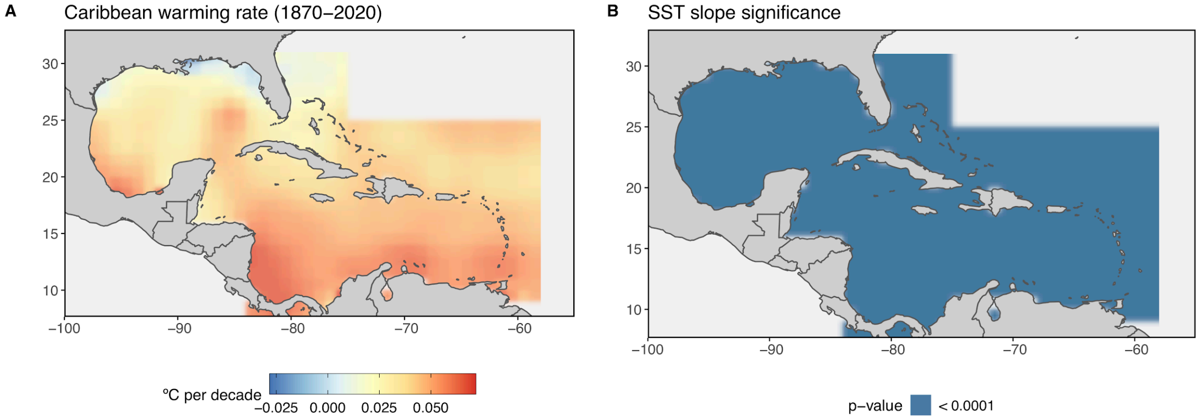


**Supplemental Figure 6**: **A)** Rate of SST change (°C per decade) over the duration of the HadISST database across the Caribbean from 1871 to 2020 (mean slope 0.04 ± 0.014 °C per decade) and **B)** significance of rate of SST change. Grey ocean area was not included in these analyses.

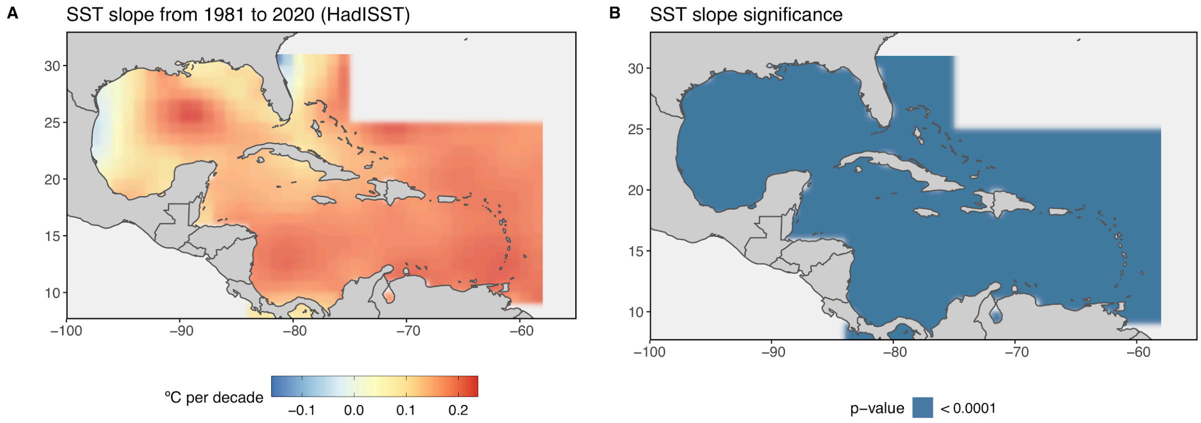


**Supplemental Figure 7**: **A)** Rate of SST change (°C per decade) across the Caribbean from 1981 to 2020 (HadISST; mean slope 0.16 ± 0.054 °C per decade) and **B)** significance of rate of SST change. Grey ocean area was not included in these analyses.

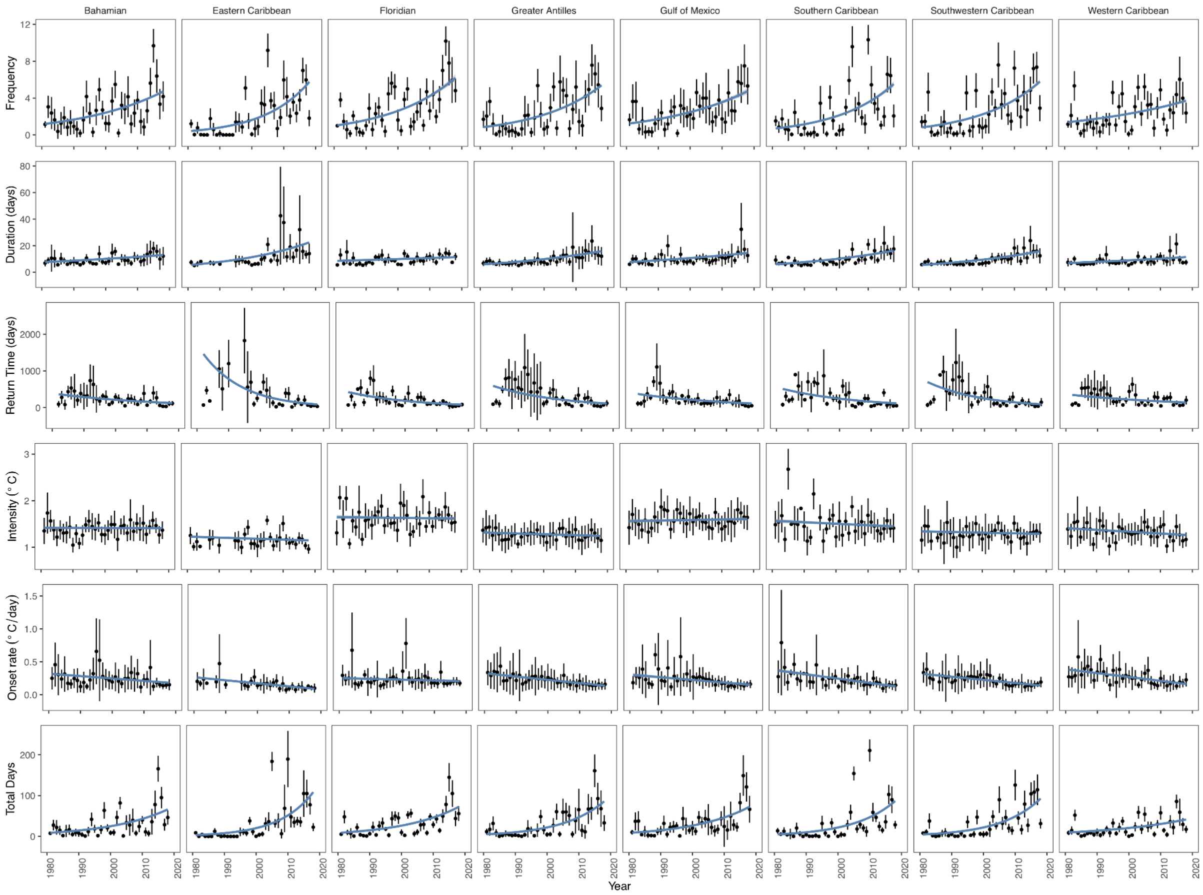


**Supplemental Figure 8**: MHW trends (1981–2018) across Caribbean coral reefs by ecoregion. Temperature data are based on OISST gridded data to determine frequency (number events per year), duration (number days per event), return time (number days per event) since the previous event, onset rate (°C per day) from start until peak intensity, peak intensity (°C), and total days reefs experience MHWs per year. Points denote annual mean values (±SD) and blue lines represent linear (lm or glm) trends within each ecoregion (see **Figures 1, Supplemental Figure 4** for ecoregion locations). Frequency, duration, and return time across all Caribbean coral reefs are depicted in **Figure 5** in the main text.

### Supplemental Tables

**Supplemental Table 1.** Marine heatwave (MHW) properties examined in this study, as developed by Hobday et al. 2016[1].

| Metric | Units | Description |
|---|---|---|
| Frequency | Number of events | Number of discrete MHW events |
| Total MHW days | days | The total number of days a location experienced a MHW per year |
| Duration | days | The number of days between the start and end date of each distinct MHW event |
| Peak intensity | °C | The maximum temperature, above the seasonal varying climatological mean, reached during the MHW event |
| Onset rate | °C/day | The rate of change in temperature between the event start date and date of peak intensity |
| Return time | days | The number of days elapsed since a previous MHW event in that location |

**Supplemental Table 2.** Mean ocean warming rate (°C per decade; with 95% confidence interval) and total increase in temperature (°C) on coral reefs within each Caribbean ecoregion since the noted year of inflection point. The inflection point for each ecoregion was identified as the year in which annual warming rates significantly increased based on the first derivative of the GAM curve for each ecoregion (**see Figures 3 and Supplemental Figure 4**).

| **Ecoregion** | **Inflection point** | **Mean rate (°C per decade)** | **Warming extent (°C)** |
|---|---|---|---|
| Bahamian | 1988 | 0.17 *(0.161-0.182)* | 0.53 |
| Eastern Caribbean | 1984 | 0.26 *(0.260-0.267)* | 0.91 |
| Floridian | 1993 | 0.22 *(0.207-0.231)* | 0.57 |
| Greater Antilles | 1986 | 0.16 *(0.160-0.168)* | 0.53 |
| Gulf of Mexico | 1981 | 0.21 *(0.186-0.241)* | 0.80 |
| Southern Caribbean | 1981 | 0.26 *(0.258-0.268)* | 0.99 |
| Southwestern Caribbean | 1982 | 0.20 *(0.194-0.204)* | 0.74 |
| Western Caribbean | 1999 | 0.24 *(0.235-0.253)* | 0.48 |

**Supplemental Table 3**. Model results for MHW trends on coral reefs in the Caribbean basin between 1981-2018. Frequency was modeled using a glm model with poisson distribution and log link. Estimates, standard error, z scores (statistic), and p-values are reported. Nagelkerke pseudo r-squared used to assess goodness of fit. MHW duration and return time were log transformed before modeling with ols models. Estimate, standard error, t value (statistic) and p-value are reported along with multiple and adjusted r-squared.

| | **Estimate** | **Standard error** | **Statistic** | **P-value** |
|---|---|---|---|---|
| **Frequency** | | | | |
| *Intercept* | -93.15 | 0.892 | -104.4 | < 0.001 |
| Year | 0.05 | 0 | 105.6 | < 0.001 |
| $R^2$ | 0.4800 | | | |
| **Duration** | | | | |
| *Intercept* | -36.51 | 0.666 | -54.8 | < 0.001 |
| Year | 0.02 | 0 | 58.2 | < 0.001 |
| $R^2$ | 0.1850 | | | |
| **Return Time** | | | | |
| *Intercept* | 87.97 | 1.668 | 52.8 | < 0.001 |
| Year | -0.04 | 0.001 | -49.8 | < 0.001 |
| $R^2$ | 0.1480 | | | |

**Supplemental Table 4.** Mean decadal value of MHW parameters (frequency, duration, and return time) for the entire basin (Caribbean) and by ecoregion.

| *Duration (days)* | 1980s | 1990s | 2000s | 2010s |
|---|---|---|---|---|
| **Caribbean** | **7.6** | **8.3** | **10.8** | **14.5** |
| Bahamian | 8.6 | 8.6 | 10.3 | 12.8 |
| Eastern Caribbean | 6.9 | 8.1 | 15.0 | 19.0 |
| Floridian | 8.9 | 9.5 | 9.6 | 11.3 |
| Greater Antilles | 7.3 | 7.9 | 11.1 | 14.4 |
| Gulf of Mexico | 8.2 | 10.8 | 9.5 | 14.6 |
| Southern Caribbean | 7.5 | 7.5 | 10.3 | 15.7 |
| Southwestern Caribbean | 7.3 | 7.5 | 9.6 | 15.3 |
| Western Caribbean | 7.1 | 8.4 | 8.9 | 11.4 |
| *Frequency (events per year)* | | | | |
| **Caribbean** | **1.2** | **1.6** | **2.8** | **4.3** |
| Bahamian | 1.6 | 2.0 | 2.6 | 4.2 |
| Eastern Caribbean | 0.5 | 1.0 | 2.7 | 4.1 |
| Floridian | 1.2 | 2.7 | 2.5 | 5.4 |
| Greater Antilles | 1.2 | 1.5 | 2.9 | 4.4 |
| Gulf of Mexico | 1.5 | 2.0 | 3.0 | 4.1 |
| Southern Caribbean | 1.0 | 1.2 | 2.9 | 4.5 |
| Southwestern Caribbean | 1.2 | 1.4 | 3.1 | 4.7 |
| Western Caribbean | 1.6 | 2.0 | 2.7 | 3.2 |
| *Return Time (days)* | | | | |
| **Caribbean** | **376.7** | **373.2** | **202.4** | **110.8** |
| Bahamian | 304.6 | 264.9 | 201.3 | 122.2 |
| Eastern Caribbean | 721.9 | 651.1 | 236.3 | 94.9 |
| Floridian | 334.2 | 234.7 | 182.0 | 83.1 |
| Greater Antilles | 398.9 | 435.0 | 202.1 | 107.8 |
| Gulf of Mexico | 364.9 | 247.7 | 158.2 | 133.6 |
| Southern Caribbean | 327.8 | 442.4 | 202.6 | 106.4 |
| Southwestern Caribbean | 449.2 | 422.2 | 194.1 | 96.2 |
| Western Caribbean | 299.3 | 257.8 | 204.7 | 145.9 |

### References

1. Habary, A., Johansen, J. L., Nay, T. J., Steffensen, J. F. & Rummer, J. L. Adapt, move or die – how will tropical coral reef fishes cope with ocean warming? *Global Change Biology* **23**, 566–577 (2017).
